## Supplementary Information for "Social stratification without genetic differentiation at the site of Kulubnarti in Christian Period Nubia"

Sirak et al.

#### Table of Contents

**Supplementary Figures……………………………………………………..………..………..…2**

**Supplementary Tables……….……………………………...…………………………...…8**

**Supplementary Notes**

**Supplementary Note 1. Background of Kulubnarti……….……………………......11**

**Supplementary Note 2. Radiocarbon dating……………….………………........……18**

**Supplementary Note 3. Genetic relatedness and consanguinity...…...…………..…...21**

**Supplementary Note 4. *qpAdm*…………………………………..…………………..24**

**Supplementary Note 5. Mitochondrial DNA analysis and haplogroup calling……...26**

**Supplementary References……………………………………………………...…...……33**

### Supplementary Figures

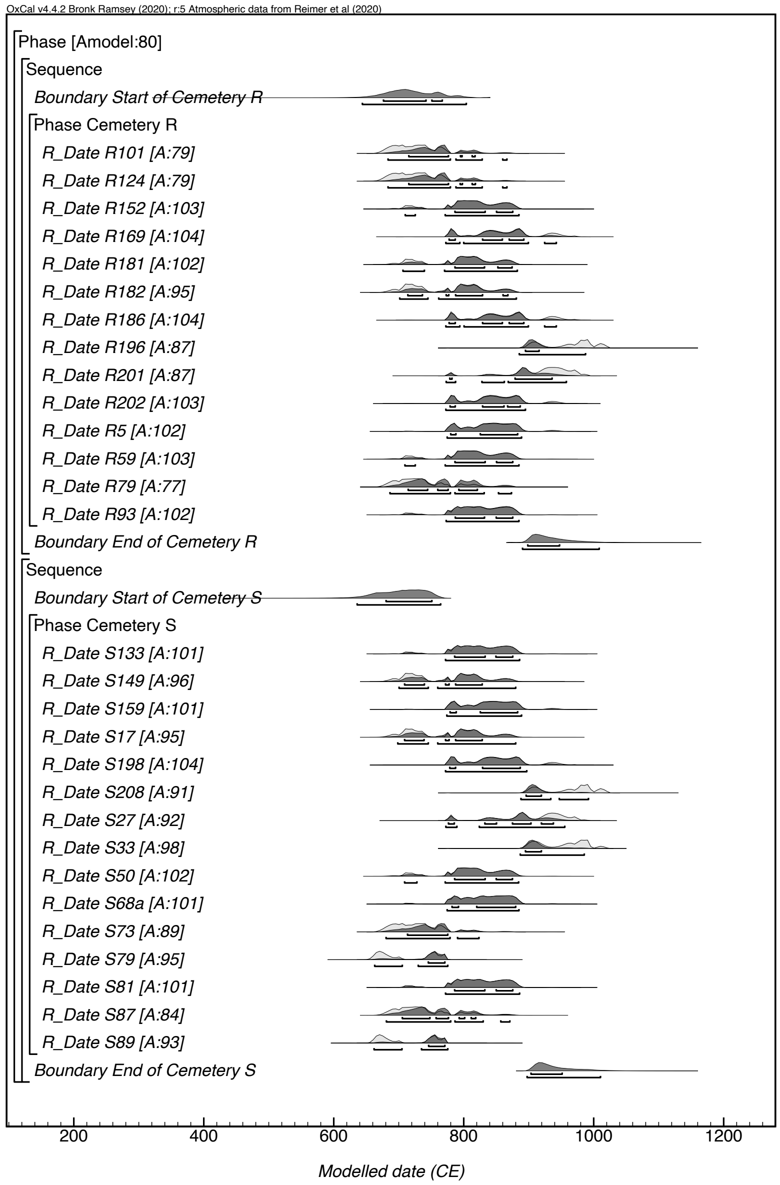


**Supplementary Figure 1.** Radiocarbon ($^{14}C$) dates for 29 individuals from Kulubnarti (14 from the R cemetery and 15 from the S cemetery). Labels are consistent with 'Skeletal Code' as in Supplementary Data 2. Details of $^{14}C$ dating are in Supplementary Note 2.

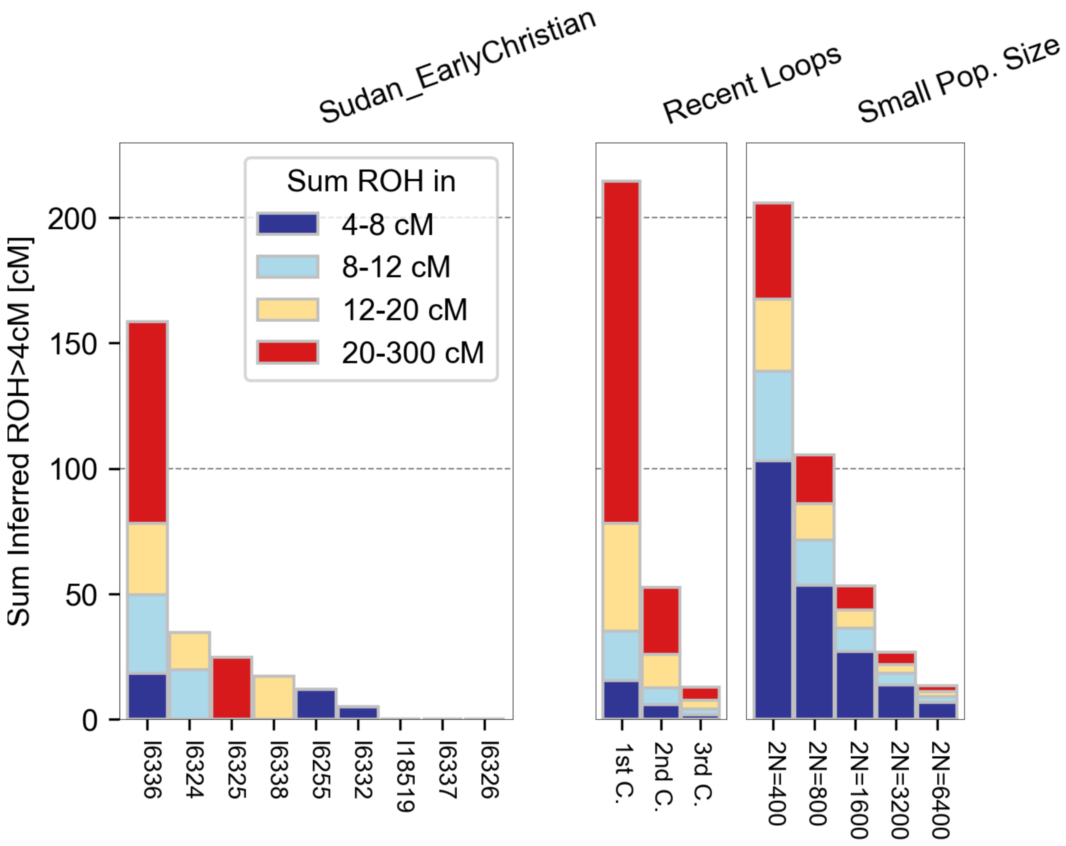


**Supplementary Figure 2. ROH calls in ancient individuals**. We depict inferred ROH for 9 ancient individuals with >400,000 SNPs; labels are consistent with 'Master ID (Lab)' in Supplementary Data 2. Each vertical bar represents one individual, and we depict the total sums of ROH that fall into four length categories: 4-8cM (dark blue), 8-12cM (light blue), 12-20cM (yellow), and >20cM (red) for each individual. 'Recent Loops' and 'Small Pop. Size' show analytical expectations and are calculated using the formulas reported in ref.[1]. "C." represents "cousin."

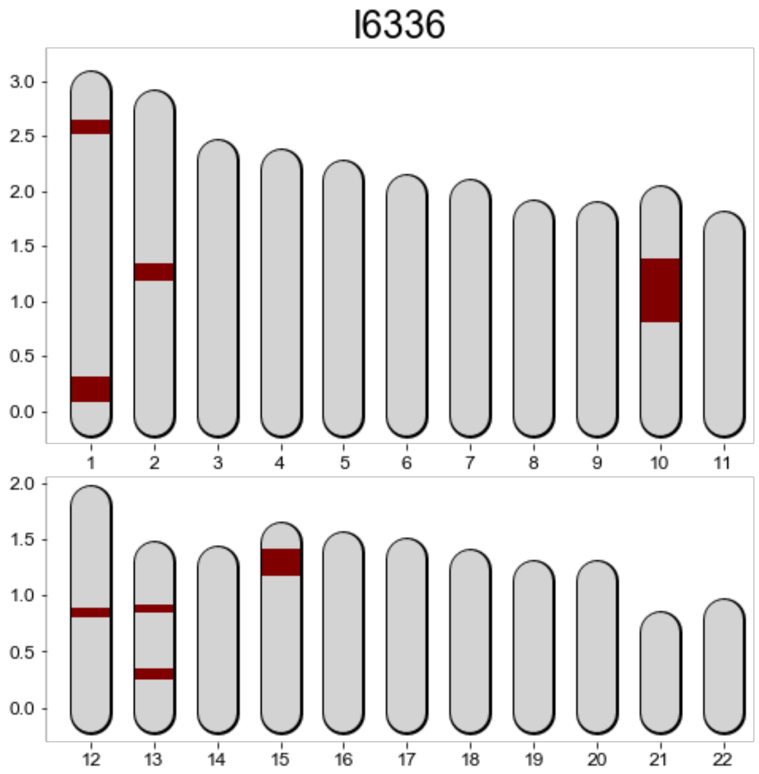


**Supplementary Figure 3. Illustration of ROH blocks >4 cM for individual I6336/S27.** This individual has a total of 158.5cM of his genome in ROH blocks >4cM, with 20cM blocks comprising over half of the total sum (~80.2cM). We identified one extremely long block of ROH (~60cM) spanning a substantial part of Chromosome 10. The number and length of ROH in I6336 provides evidence of his parents being first cousins or a close genetic equivalent (see also Supplementary Figure 2). The units on the y-axis denote the sex-averaged genomic map length measured in Morgan.

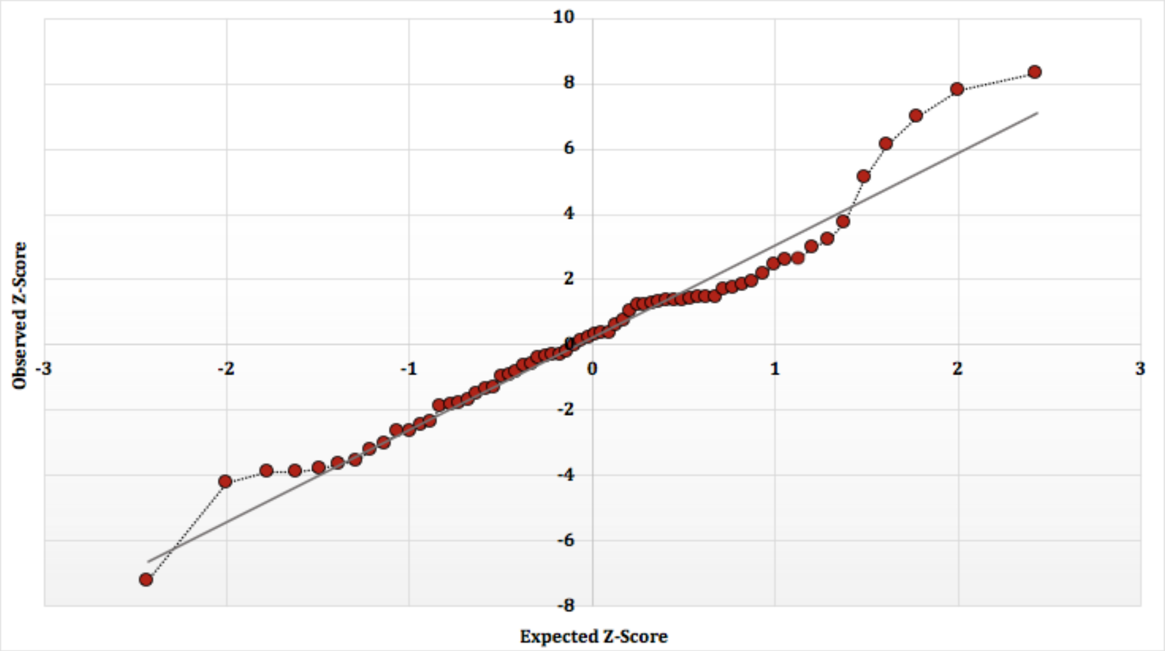


**Supplementary Figure 4.** Q-Q plot showing deviation from normal distribution of Z-scores for $f_4$(*Nilotic_Test*, *WestEurasia_Test*; *Individual*, *Kulubnarti_Without_Individual*). Data from Supplementary Data 6. Only ~27% of individuals fall within one standard deviation of the mean, while ~62% fall within two standard deviations, and ~76% fall within three standard deviations.

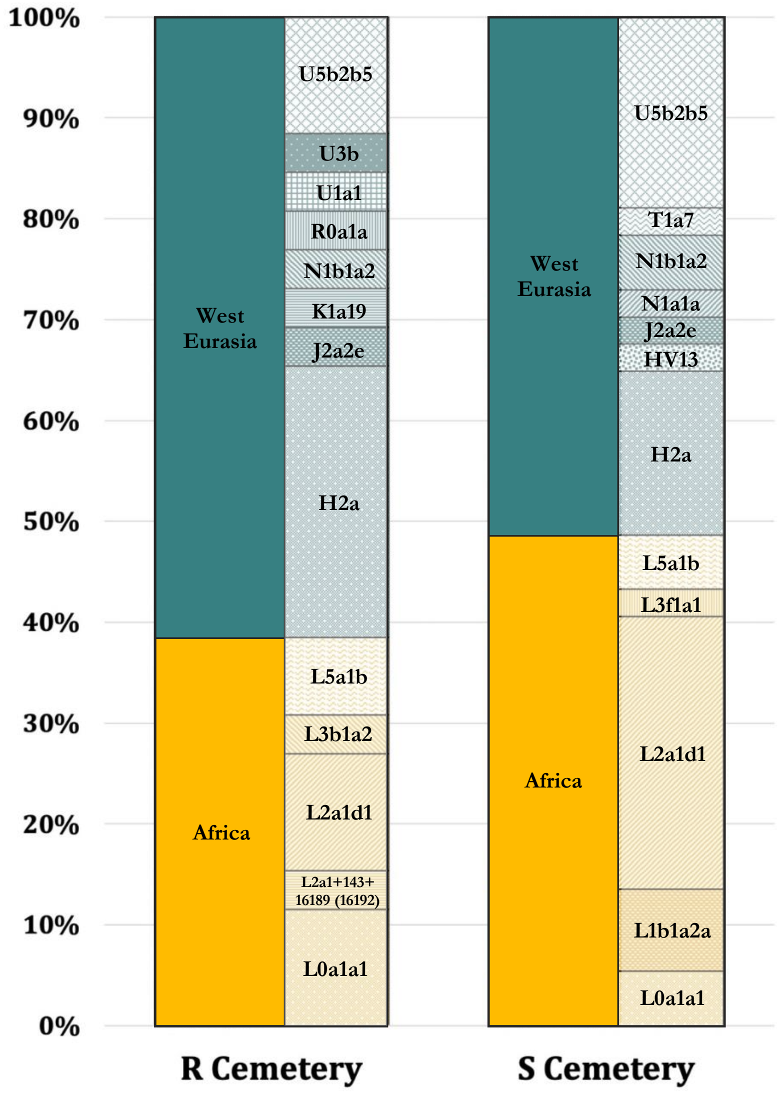


**Supplementary Figure 5.** mtDNA haplogroup calls for 63 individuals who were not first-degree relatives sharing a maternal lineage divided by cemetery of burial and grouped by most likely geographic region of origin and primary geographic distribution; data are in Supplementary Data 12.

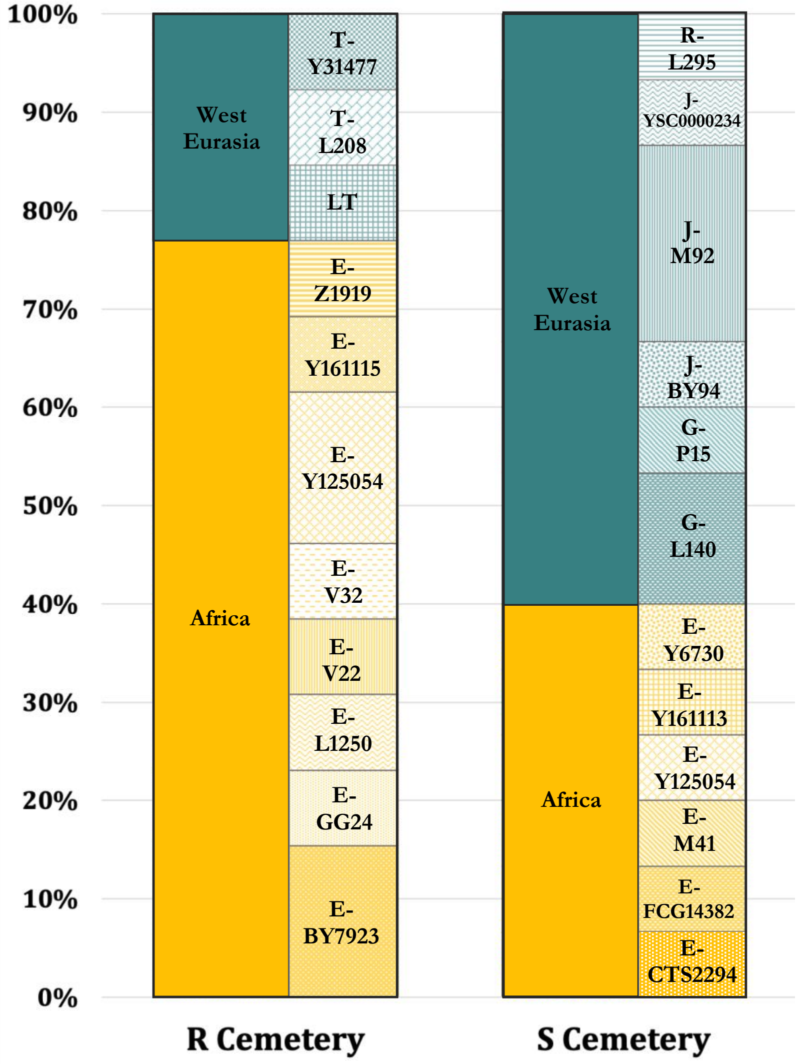


**Supplementary Figure 6.** Y haplogroup calls in terminal mutation notation for 28 males divided by cemetery of burial who were not first-degree relatives and grouped by most likely geographic region of origin and primary geographic distribution; data are in Supplementary Data 13.

### Supplementary Tables

#### Supplementary Table 1. Radiocarbon dates for 29 individuals from Kulubnarti.

| UGAMS# | Master ID (Lab) | Skeletal code | Collagen yield, % | Atomic C:N ratio | $\delta^{13}C$, ‰ VPDB | $^{14}C$ age years, BP | ± | Unmodeled date, cal CE (68.3% probability range) | Unmodeled date, cal CE (95.4% probability range) |
|---|---|---|---|---|---|---|---|---|---|
| 34382 | I6138 | R101 | 15.9 | 3.4 | -16.5 | 1270 | 22 | 680–770 | 660–820 |
| 34383 | I6139 | R124 | 16.5 | 3.3 | -17.0 | 1270 | 22 | 680–770 | 660–820 |
| 34384 | I6251 | R152 | 16.1 | 3.7 | -17.2 | 1220 | 22 | 780–880 | 700–890 |
| 34385 | I6340 | R169 | 12.3 | 3.5 | -18.3 | 1170 | 22 | 770–940 | 770–960 |
| 34386 | I6252 | R181 | 15.4 | 3.5 | -15.7 | 1230 | 22 | 700–880 | 700–890 |
| 34387 | I6140 | R182 | 17.6 | 3.4 | -16.7 | 1240 | 22 | 700–830 | 680–880 |
| 34388 | I6141 | R186 | 17.2 | 3.3 | -17.2 | 1170 | 22 | 770–940 | 770–960 |
| 34389 | I6327 | R196 | 15.6 | 3.8 | -17.6 | 1080 | 23 | 890–1020 | 890–1030 |
| 34390 | I6328 | R201 | 16.7 | 3.5 | -16.4 | 1140 | 22 | 880–980 | 770–990 |
| 34391 | I6253 | R202 | 15.9 | 3.6 | -18.3 | 1180 | 22 | 770–890 | 770–950 |
| 34392 | I6329 | R5 | 15.9 | 3.5 | -17.5 | 1190 | 22 | 770–890 | 770–900 |
| 34393 | I6250 | R59 | 16.9 | 3.5 | -17.3 | 1220 | 22 | 780–880 | 700–890 |
| 34394 | I6330 | R79 | 16.0 | 3.3 | -17.0 | 1260 | 22 | 680–770 | 670–830 |
| 34395 | I6331 | R93 | 15.5 | 3.5 | -17.8 | 1210 | 22 | 780–880 | 700–890 |
| 34396 | I6324 | S133 | 15.1 | 3.4 | -17.0 | 1210 | 22 | 780–880 | 700–890 |
| 34397 | I6258 | S149 | 6.3 | 3.9 | -19.1 | 1240 | 22 | 700–830 | 680–880 |
| 34398 | I6332 | S159 | 13.2 | 3.4 | -17.8 | 1190 | 22 | 770–890 | 770–900 |
| 34399 | I6333 | S17 | 15.1 | 3.4 | -18.0 | 1240 | 22 | 700–830 | 680–880 |
| 34400 | I6334 | S198 | 18.3 | 3.8 | -17.5 | 1180 | 24 | 770–890 | 770–950 |
| 34401 | I18519 | S208 | 18.4 | 4.8 | -18.1 | 1080 | 22 | 890–1020 | 890–1030 |
| 34402 | I6336 | S27 | 11.9 | 3.7 | -17.8 | 1150 | 23 | 770–980 | 770–980 |
| 34403 | I6254 | S33 | 14.3 | 3.4 | -17.2 | 1090 | 22 | 890–1000 | 890–1020 |
| 34404 | I6255 | S50 | 17.6 | 3.4 | -16.2 | 1220 | 22 | 780–880 | 700–890 |
| 34405 | I6256 | S68a | 12.7 | 3.4 | -16.3 | 1200 | 22 | 780–880 | 770–890 |
| 34406 | I6325 | S73 | 13.2 | 3.4 | -17.9 | 1270 | 22 | 680–770 | 660–820 |
| 34407 | I6337 | S79 | 24.1 | 3.3 | -15.9 | 1320 | 22 | 660–780 | 650–780 |
| 34408 | I6257 | S81 | 2.8 | 3.6 | -16.3 | 1210 | 22 | 780–880 | 700–890 |
| 34409 | I6326 | S87 | 11.0 | 3.5 | -16.9 | 1260 | 22 | 680–770 | 670–830 |
| 35229 | I6338 | S89 | 14.1 | 3.3 | -17.1 | 1320 | 20 | 660–780 | 650–780 |

**Supplementary Table 2. Families at Kulubnarti.** Inter-cemetery kin pairs in bold; relationship (if known) included. Relatives called following the method in ref.[2].

| Family | Relative 1 | Relative 2 | Intra- or inter-cemetery | Relationship (Degree) | Details |
|---|---|---|---|---|---|
| A | I18522/S235 | I18521/S21 | intra-cemetery | 3rd-4th | |
| A | I18507/S114 | I18522/S235 | intra-cemetery | 2nd | |
| A | I18507/S114 | I17449/S147 | intra-cemetery | 3rd-4th | |
| A | I17449/S147 | I18522/S235 | intra-cemetery | 2nd | |
| **B** | **I17450/S51** | **I19015/R21** | **inter-cemetery** | **2nd** | |
| C | I19145/R173 | I6330/R79 | intra-cemetery | 2nd | |
| D | I6256/S68a | I17475/S144 | intra-cemetery | 1st | siblings |
| E | I6138/R101 | I6331/R93 | intra-cemetery | 1st | brothers |
| E | I6331/R93 | I19143/R150 | intra-cemetery | 3rd-4th | |
| **E** | **I6337/S79** | **I19143/R150** | **inter-cemetery** | **3rd-4th** | |
| F | I6250/R59 | I6251/R152 | intra-cemetery | 2nd | |
| **F** | **I6251/R152** | **I6255/S50** | **inter-cemetery** | **3rd-4th** | |
| F | I6255/S50 | I6333/S17 | intra-cemetery | 2nd | |
| F | I6333/S17 | I18514/S182 | intra-cemetery | 3rd-4th | |
| G | I18612/S16 | I18610/S29 | intra-cemetery | 1st | father (I18612/S16) and son (I18610/S29) |
| G | I6336/S27 | I18610/S29 | intra-cemetery | 3rd-4th | |
| G | I6324/S133 | I6336/S27 | intra-cemetery | 3rd-4th | |
| G | I6336/S27 | I18518/S201 | intra-cemetery | 3rd-4th | |
| G | I6336/S27 | I17481/S45 | intra-cemetery | 3rd-4th | |
| G | I18525/S37 | I6336/S27 | intra-cemetery | 2nd | |
| G | I6324/S133 | I18509/S132 | intra-cemetery | 1st | sisters |
| G | I18538/S53 | I6324/S133 | intra-cemetery | 2nd | |
| G | I18538/S53 | I18509/S132 | intra-cemetery | 3rd-4th | |
| **H** | **I6325/S73** | **I19132/R57** | **inter-cemetery** | **3rd-4th** | |
| **H** | **I19134/R84** | **I6254/S33** | **inter-cemetery** | **2nd** | |
| **H** | **I6254/S33** | **I19132/R57** | **inter-cemetery** | **3rd-4th** | |
| **H** | **I17451/S15** | **I19132/R57** | **inter-cemetery** | **uncertain** | **low SNP coverage precludes us from determining the exact degree of this relationship** |
| H | I19132/R57 | I19134/R84 | intra-cemetery | uncertain | low SNP coverage precludes us from determining the exact degree of this relationship |

**Supplementary Table 3. Runs of homozygosity (ROH) segments >4cM for individuals with sufficient coverage (>400,000 SNPs covered).** Number of ROH segments longer than a specified size denoted by 'n_roh' and sum of ROH segment lengths longer than a specified size denoted by 'sum_roh.'

| Master ID (Lab) | Skeletal Code | sum_roh >4 | n_roh >4 | sum_roh >8 | n_roh >8 | sum_roh >12 | n_roh >12 | sum_roh> 20 | n_roh >20 |
|---|---|---|---|---|---|---|---|---|---|
| I18519 | S208 | 0 | 0 | 0 | 0 | 0 | 0 | 0 | 0 |
| I6337 | S79 | 0 | 0 | 0 | 0 | 0 | 0 | 0 | 0 |
| I6326 | S87 | 0 | 0 | 0 | 0 | 0 | 0 | 0 | 0 |
| I6332 | S159 | 5.1613 | 1 | 0 | 0 | 0 | 0 | 0 | 0 |
| I6255 | S50 | 12.0226 | 2 | 0 | 0 | 0 | 0 | 0 | 0 |
| I6338 | S89 | 17.3308 | 1 | 17.3308 | 1 | 17.3308 | 1 | 0 | 0 |
| I6325 | S73 | 24.7553 | 1 | 24.7553 | 1 | 24.7553 | 1 | 24.7553 | 1 |
| I6324 | S133 | 34.6752 | 3 | 34.6752 | 3 | 14.8383 | 1 | 0 | 0 |
| I6336 | S27 | 158.5166 | 10 | 139.9770 | 7 | 108.9109 | 4 | 80.2499 | 2 |

**Supplementary Table 4. FST between the individuals in the Kulubnarti R and S cemeteries.**

| A | B | $F_{ST}$ | std err |
|---|---|---|---|
| Kulubnarti_R | Kulubnarti_S | 0.001331 | 0.00053 |

### Supplementary Note 1 – Background of Kulubnarti

The site of Kulubnarti ("Island of Kulb" in the Mahasi dialect of Nubian) is located on the bank of the Nile River in Sudanese Nubia, approximately 130 kilometers south of the present-day border between Egypt to the north and Sudan to the south. The site was first discovered in 1969 as part of the United Nations Educational, Scientific, and Cultural Organization (UNESCO) International Campaign to Save the Monuments of Nubia that aimed to identify and excavate archaeological sites that would be inundated by the creation of a permanent lake (Lake Nasser/Lake Nubia) following the construction of the High Aswan Dam.

We consider Kulubnarti to include both the large trapezoidal island of Kulb (dimensions approximately 1 km by 2 km) as well the adjacent west bank. Prior to the creation of Lake Nubia, Kulubnarti was a headland projecting into the Nile River from the western bank. While the island and mainland are presently separated by a narrow channel, they were connected during the Christian Period (~550-1400 CE) except at the peak of the Nile flood[3]. Excavation of the village of Kulubnarti and survey of the associated cemeteries occurred in 1969. Excavation of these two contemporaneous and geographically-proximate cemeteries, one on Kulubnarti island and one on the western bank opposite the south end of the island, occurred in 1979 under a license granted by the Sudan Government Antiquities Service to Dr. William Y. Adams at the University of Kentucky and funded by the National Science Foundation (Grant No. 77-270210-535); the work was undertaken as a contribution to the International Campaign to Save the Monuments of Nubia[3]. This campaign was conducted in response to the impending destruction of Nubia caused by the construction of a new High Aswan Dam, which started in 1947. Unlike the reservoir created by the first Aswan Dam, which was emptied during part of each year, the High Aswan Dam created a permanent lake. More than forty expeditions were consequently planned and conducted between the Egyptian border and the head of the proposed reservoir prior to the Dam's construction. These expeditions discovered over 1,000 archaeological sites and excavated nearly one-third of them[4]. The 1979 excavation of the Kulubnarti cemeteries was conducted as a Joint Colorado-Kentucky Expedition, led by Dr. Dennis van Gerven (co-senior author on this manuscript). Over one field season lasting two months, the remains of 406 individuals were recovered from two cemeteries and were sent to the University of Colorado at Boulder (UC Boulder; USA) for analysis and curation; this osteological material is currently curated at Arizona State University (ASU; USA).

Detailed information regarding the architectural[3], artifactual[5], and human[6] remains from Kulubnarti have been published in a series of three monographs. An overview of the site, the cemeteries, and the people who inhabited Kulubnarti during the Christian Period will be provided here, and the genetic data newly generated by this work will be presented in the main manuscript and integrated into the existing framework built by archaeological work.

**The site of Kulubnarti**

Kulubnarti is situated within the *Batn el Hajar* region of Nubia, an inhospitable area that separated Egyptian-influenced Lower Nubia (the land between the First and Second Cataracts of the Nile) and the rest of Upper Nubia (the land below the Second Cataract of the Nile)[7] (see Figs. 1a and 1b in the main manuscript). This region is described to be the most "barren and forbidding of all Nubian environments," with a "lunar" feeling characterized by steep riverbanks, countless *jebels* (large outcroppings of rock), and sharp *wadis* (channels that are dry except in the rainy season)[4,8]. The Nile becomes unnavigable in the *Batn el Hajar,* coursing through granite rapids and hundreds of riverine islands. While this area supported scattered populations who built small villages and hamlets clustered around the region's few floodplains, this terrain also functioned as a natural deterrent against the infiltration by foreign peoples. Based on a combination of environmental and political factors, the *Batn el Hajar* garnered the reputation as being the "granite curtain" that dissuaded the movement of peoples from Egypt and the Arabic world southward along the Nile corridor[4]. It is specifically suggested that this "granite curtain" protected the inhabitants of Upper Nubia from the gradual expansion of Islamic influence until at least the 12$^{th}$ century CE[4].

Kulubnarti was a small hamlet located on the bank of the Nile at a location where some alluvial soil was present, but also where there was no continuous floodplain. Archaeological evidence suggests that it is likely that population density at Kulubnarti (consistent with other sites in the *Batn al Hajar*) was consistently low, and that the population that inhabited this site was always relatively impoverished compared to contemporaneous populations in more fertile regions along the Nile River to the north and south[9]. While there is some evidence of mat, basket, and sandal-making at Kulubnarti throughout the entirety of the Christian Period, a notable lack of specialized craft and imported goods suggests that subsistence agriculture was likely the main activity[5,9].

Due to a paucity of arable land, individual landholdings at Kulubnarti were very small and highly-valued. The channel that separates the island from the mainland was farmed using *seluka* cultivation (a type of cultivation practice that relies on alluvium gradually exposed as the Nile receded from its annual flood) to grow legumes and other fodder crops[3,4,6,10]. Isotopic data suggest that dietary consumption at Kulubnarti was primarily based on 'winter' $C^3$ plants harvested in April (including barley, legumes, and wheat), with some consumption of 'summer' $C^4$ plants harvested in June (including sorghum and millet)[11-13]; this is largely consistent with dietary patterns throughout rural areas of Nubia today[4,10]. Animals (including goats, cattle, sheep, and pigs) were kept in small numbers, but animal meat was uncommon in the diet[9,11,13]; instead the Nubians obtained their protein primarily from plant sources[4,14]. The range of isotopic values and lack of archaeological evidence suggests that consumption of riverine products such as fish was rare, indicating that the Kulubnarti Nubians subsisted upon a terrestrially-based diet[4,11].

**The Kulubnarti cemeteries**

Human skeletal remains were recovered from two cemeteries at Kulubnarti. Site 21-S-46 (the 'S cemetery') was situated within a dry ancient *wadi* near the west side of Kulubnarti Island, and site 21-R-2 (the 'R cemetery') was located on the mainland's west bank[6] (see Fig. 1b).

In the S cemetery, the earliest graves were of pre-Christian type, with a clear transition into Christian-style graves, identified by grave orientation, body positioning, and burial shrouds[6]. The total number of graves in the S cemetery remains unknown, though estimations place this number at approximately 300[6]. During the 1979 excavation, 218 graves were excavated, and 215 bodies were uncovered from this cemetery. Most of the graves were slot graves (straight-sided pits with rounded or square ends) that had a covering at the surface, most often simple pavements of flat but unshaped granite slabs arranged in a rectangle over the top of the grave[6]. Each of the excavated graves had an east-west orientation with the head of the body placed at the west end, as is typical for Christian-style burials[6]. Most individuals, regardless of sex or age, were wrapped in a shroud; however, as is common with Christian burials, recovery of any personal goods included in the graves was rare[6].

The R cemetery was located next to a Classic Christian Period (850–1100 CE) domed church as well as an Early Christian Period (~550-800 CE) walled settlement[6]. In addition to Christian Period graves dug into a barren alluvial surface that merged with the Nile floodplain, Islamic-type graves were found at one end of the R cemetery. It was estimated that the R cemetery contained between 500 and 600 graves[6]. A total of 188 graves were opened during the 1979 expedition, all but six of which were concentrated in one contiguous area at the far western end of the cemetery. The concentration of all burials excavated from the R cemetery at the far western end of the cemetery raises the concern that this sample in particular may not be representative of the cemetery as a whole[6]; however, there is little additional evidence to suggest that it is not. From these 188 graves, a total of 191 bodies were recovered[6]. It has been noted that the graves from the R cemetery exhibited no typological distinction from those at the S cemetery[6]. While the orientation of graves at the R cemetery was more erratic than at the S cemetery, this difference has been attributed to the lack of any topographic feature on the western horizon that could serve as an orientation point. Consistent with the S cemetery, the R cemetery most frequently exhibited slot graves, high frequency of burial shrouds, and limited grave goods[6].

A lack of distinctive grave goods presented challenges for the precise dating of the Kulubnarti cemeteries[6]. The original interpretation of the available archaeological data was that the two cemeteries were used in successive periods with partial overlap. Specifically, analysis of pottery within the graves as well as architectural associations originally suggested that the S cemetery represented a population from the Early Christian Period (550–800 CE), while the presence of vaulted brick tombs and both Christian and Muslim burial styles suggested that the R cemetery was in use from the Early Christian through the Terminal Christian Period (550–1400 CE)[9,15]. Analysis of the textiles found in the graves of both cemeteries, however, suggested that the textiles found exhibited characteristics of Nubian textiles from the Early Christian Period, including a high percentage of woolen fabrics, a low percentage of cotton fabrics, an even lower percentage of flax, absence of silk, and rare occurrence of dyed color[6]. The contemporaneity of the cemeteries was further supported by a small sample of radiocarbon dates[16]; additional support for their contemporaneity is provided by the 29 new radiocarbon dates assembled as part of this work (see Supplementary Table 1 for direct dates).

**Human osteological remains from Kulubnarti**

Following their excavation and exportation to UC Boulder, an assessment of age and sex was conducted by Dr. Dennis Van Gerven. Developmental age at death was determined based on a seriation technique that examined inter-individual variation in multiple well-established ageing criteria, including stages of dental eruption[17,18], epiphyseal fusion[18,19], and age-related changes in the pubic bones[20-22]. Population-specific patterns of dental attrition and skeletal degenerative changes were also considered[6]. Age estimations were made for 399 out of 406 individuals by arranging all individuals in a graded developmental series. Sex was determined for adults based on dimorphic skeletal features, including features of the pelvis[23,24], cranium[19,25], and long bones[26]. Residual soft tissue occasionally enabled the determination of sex in subadults. All data are published in ref.[6]. Both morphological and genetic sex (determined as part of this work) are provided in Supplementary Data 1.

Most individuals from Kulubnarti appeared macroscopically well-preserved because the heat and aridity of the Nubian environment encouraged soft tissue preservation; over half of the individuals recovered had some preserved soft tissue in the form of skin, tendons, or muscles, and over one-third of the individuals still had hair. In addition to heat, the soft tissue preservation is due in part to the location of both Kulubnarti cemeteries away from the Nile flood. Specifically, though the Nile flooded annually, the location of the S cemetery was at least 10 meters above the level of the Nile floodplain in the Early Christian Period and had not been exposed to flooding in recent millennia. The R cemetery sample was selected from the area of highest ground in the cemetery, making it unlikely that any burials were affected by flooding[6].

Several decades of bioarchaeological research were conducted on the human osteological remains from Kulubnarti. Two primary observations were made through this work. The first observation was that studies of biological distance ('biodistance') suggested a close biological relationship between the individuals recovered from the Kulubnarti R and S cemeteries. Comparing craniometric data collected from individuals recovered from the R and S cemeteries to a time-series from Wadi Halfa (located ~130km to the north), Van Gerven[27] determined that the principal discrimination was between Wadi Halfa and Kulubnarti, while the least significant difference was between the R and S cemeteries. Morphological similarity between individuals in the R and S cemeteries was also detected through the analysis of cranial nonmetric traits[28]. In addition, the application of multivariate statistics to discrete dental data identified no significant variation between the R and S cemeteries[29].

Despite morphological similarity, the second observation was that individuals buried in the S cemetery were exposed to more stress, experienced more ill-health, and died younger than the individuals buried in the R cemetery. Cribra orbitalia, a commonly-used indicator of generalized stress, was found in 94% of S cemetery children in comparison to 82% of R cemetery children, indicating a higher degree of childhood stress for individuals buried in the S cemetery[15,30]. A similar pattern was seen for linear enamel hypoplasias (LEHs), another indicator of generalized stress. While a nearly universal presence of LEH lesions were found in both cemeteries at Kulubnarti, the lesions appeared more frequently and were maintained at a higher frequency for longer in individuals from the S cemetery, leading to a prolonged period of intensified childhood mortality[31]. The increased frequency of stress-induced lesions found in individuals buried in the S cemetery corresponds to increased childhood mortality. Mean life expectancy computed from composite life tables[32] revealed that while differences in mortality after childhood were minimal, mortality between birth and age eight was significantly higher for individuals buried in the S cemetery than the R cemetery[6]. Therefore, probabilities of dying were not only higher for the children interred in the S cemetery, but chances of dying remain higher for longer[9]. This resulted in an average life expectancy of 10.6 years for the S cemetery overall compared to 18.8 years for the R cemetery[9]. The differences in morbidity and mortality between the S and R cemeteries were not attributable to variation in diet[11]. Analysis of carbon, nitrogen, and oxygen isotopes from bone tissue indicates no significant relationships between isotopic indicators and cemetery of burial, suggesting no isotopically-measurable differences in proportional dietary composition[11].

These two observations made using the human osteological material from Kulubnarti, taken together with material (textile) evidence of more prosperous people buried in the R cemetery than the S cemetery, but otherwise little difference in grave types or grave goods, led researchers to conclude that two biologically-related and culturally-indistinguishable, but socially-distinct groups of people lived side-by-side at Kulubnarti, and that one of these groups was considerably better-off than the other[6]. To explain this possible social stratification, anthropologists drew upon ethnographic evidence from present-day Nubia that describes groups of impoverished, landless, semi-nomadic persons who act as sharecroppers or seasonal laborers for landowning Nubian families, otherwise living off small flocks of sheep and goats. These people are known locally as the Nubian ‘underclass’[33]. The well-evidenced disparity between the people buried in the R and S cemeteries at Kulubnarti supported a hypothesis that such a social structure might have existed during the Christian Period as well, and that the individuals buried in the S cemetery were a group of itinerant and disadvantaged individuals who provided labor for the people buried in the relatively more prosperous R cemetery[33]. However, while social stratification is a common feature of complex landowning societies[34,35], the presence of a semi-nomadic, landless underclass in Christian Period Nubian society has been described as a "wholly unexpected possibility" for which there is "neither textual evidence nor archaeological evidence from other sites to support such an interpretation"[6]. In this work, we divide the individuals from the R and S cemeteries into two cemetery groups when it is necessary to carry out analyses that investigate potential differences between the people buried in each cemetery at Kulubnarti.

### Supplementary Note 2 – Radiocarbon Dating

Radiocarbon dating was performed at the Center for Applied Isotope Studies (CAIS), University of Georgia (USA) for 29 individuals that yielded genome-wide data. Collagen was extracted following the protocol in ref.[36], modified as described here. A sub-sample of bone was removed using a Dremel tool outfitted with a diamond cutting wafer. Surface contamination was removed from the sub-sample using a scalpel and wire-bristle brush; the sub-sample was simultaneously reduced to smaller fragments (approximately 3–5mm in size). These small fragments were demineralized in cold (4°C) 1N HCl for 24 hours, the acid was decanted, and demineralized fragments of bone were rinsed three times with ultrapure water (MilliQ). The bone fragments were then treated with 0.1M NaOH to dissolve and remove humic acids, followed by a series of ultrapure water rinses. Atmospheric $CO_2$ was eliminated through the rinsing of bone fragments with cold 1N HCl. The fragments were then rinsed again in ultrapure water to ~ pH 4 (slightly acidic) and heated at 80ºC for 8 hours. The solution was subsequently filtered through a glass fiber filter, isolating the total acid insoluble fraction ("collagen"), which was then freeze-dried. A ~5mg sub-sample of collagen was combusted at 575°C in an evacuated and sealed Pyrex tube in the presence of CuO, producing $CO_2$. The $CO_2$ sample was cryogenically purified from the other reaction products and catalytically converted to graphite following the method of ref.[37]. Graphite $^{14}C/^{13}C$ ratios were measured using the 0.5 MeV accelerator mass spectrometer (AMS) housed at CAIS. Sample ratios were compared to the ratio measured from the Oxalic Acid I standard (NBS SRM 4990). All results are presented as percent Modern Carbon (pMC). The quoted uncalibrated dates are given in radiocarbon years before 1950 (years BP), using a $^{14}C$ half-life of 5568 years. The date has been corrected for isotope fractionation using the $\delta^{13}C$ value measured by EA-IRMS. Uncalibrated conventional dates are presented in Supplementary Table 1.

**Bayesian chronological modeling.** Bayesian chronological modeling employed the OxCal[38] software version 4.4 and the IntCal20[39] calibration curve. We use capitalized forms of words to refer to OxCal command language terminology (i.e., Sequence, Phase, Boundary, Interval). The chronological model is expressed in terms of Sequences, Phases, and Boundaries. A Sequence is a group of events or parameters that occurred in a specific order in relation to one another. A Phase is an unordered group of events or parameters. Boundaries are used to define the boundary of a group of events, e.g., the start or end of a Phase[38]. We employed the minimum of assumptions in constructing the chronological model. The $^{14}$C data from each cemetery was modeled as an independent Phase with start and end Boundaries. We made no assumptions regarding the relative chronological relationships within or between the two Phases; each Phase is bracketed by a start and end Boundary within an independent Sequence:

```
Plot()
 {
 Curve("IntCal20"," IntCal20.14c");
 Phase()
 {
  Sequence()
  {
   Boundary("Start of Cemetery R");
   Phase("Cemetery R")
   {
    R_Date("R101", 1270, 22);
    R_Date("R124", 1270, 22);
    R_Date("R152", 1220, 22);
    R_Date("R169", 1170, 22);
    R_Date("R181", 1230, 22);
    R_Date("R182", 1240, 22);
    R_Date("R186", 1170, 22);
    R_Date("R196", 1080, 23);
    R_Date("R201", 1140, 22);
    R_Date("R202", 1180, 22);
    R_Date("R5", 1190, 22);
    R_Date("R59", 1220, 22);
    R_Date("R79", 1260, 22);
    R_Date("R93", 1210, 22);
    Interval("Duration of Cemetery R");
   };
   Boundary("End of Cemetery R");
  };
  Sequence()
  {
   Boundary("Start of Cemetery S");
   Phase("Cemetery S")
   {
    R_Date("S133", 1210, 22);
    R_Date("S149", 1240, 22);
    R_Date("S159", 1190, 22);
    R_Date("S17", 1240, 22);
    R_Date("S198", 1180, 24);
    R_Date("S208", 1080, 22);
```

```
    R_Date("S27", 1150, 23);
    R_Date("S33", 1090, 22);
    R_Date("S50", 1220, 22);
    R_Date("S68a", 1200, 22);
    R_Date("S73", 1270, 22);
    R_Date("S79", 1320, 22);
    R_Date("S81", 1210, 22);
    R_Date("S87", 1260, 22);
    R_Date("S89", 1320, 20);
    Interval("Duration of Cemetery S");
   };
   Boundary("End of Cemetery S");
  };
 };
};
```

OxCal calculates a posterior Probability Density Function (PDF) for each of these elements. We utilized the Interval command to determine the length of time in calendar years of each cemetery represented by a Phase in the model. An agreement index is calculated for each dated item ("A" values), as well as for the model as a whole ("$A_{model}$"), with A≥60 considered to be the threshold for acceptable agreement[38]. In Supplementary Figure 1, we provide individual $^{14}C$ dates. In this figure, the un-modeled calibrated date probabilities are indicated by the light gray distributions; the modeled (posterior) probabilities are shown by the dark gray distributions. The lines under the modelled distributions indicate the 68.3% highest posterior density (hpd) and 95.4% hpd ranges, the former of which are referred to in the text. In Fig. 1c, we present the modeled start and end dates (top) and duration of use (bottom) of the R and S cemeteries.

### Supplementary Note 3 – Genetic relatedness and Consanguinity

#### Genetic relatedness

We looked for genetic relatedness between all individuals in our study following the method published in ref.[2]. This method compares the mean mismatch of all autosomal SNPs with at least one sequencing read between individuals (selecting at random one read if coverage is greater than one at a particular position for a given individual). The mismatch rate is used to estimate a relatedness coefficient ($r$), which informs about the degree of relatedness between two individuals. This method is specifically applicable for estimating relatedness from haploid SNP data (common in ancient DNA analysis) and can accurately provide estimates of genetic relatedness up to third-/fourth-degree relatives.

We identify 33 individuals from Kulubnarti who share 28 genetic relationships up to the third/fourth degree (Supplementary Table 2). This included four pairs of first-degree relatives, nine pairs of second-degree relatives, 13 pairs of third-/fourth-degree relatives, and two pairs of relatives of an unknown relationship (in the latter case, the proportion of overlapping SNPs was used to infer that the individuals were related, although the low coverage precluded our ability to determine the exact degree of their relationship).

Particularly interesting in the context of two plausibly socially-stratified contemporaneous burial grounds at Kulubnarti (see Supplementary Note 1) is the identification of 7 inter-cemetery relative pairs out of the 28 total relative pairs. While comparison of the six inter-cemetery relative pairs where the degree of relatedness could be assessed against the number of relative pairs detected overall suggests that there was a modest degree of enrichment of relative pairs buried in the same cemetery versus in different cemeteries (see Table 1 in main manuscript), we were surprised to identify any cross-cemetery relatives based on the hypothesis of a caste-like social system at Kulubnarti that would have restricted inter-group mating. Instead, we find that there was more likely to be fluidity between groups and, as such, no strict caste-like system of social division. While genetic data allows us to assess the degree of biological relatedness between two individuals, it does not enable us to ascribe the social concept of "kin" onto this assessment; as such, we cannot speak to the social relationship between any two individuals (whether buried in the same cemetery or in different cemeteries). However, with no archaeological evidence of cross-cemetery relative pairs at Kulubnarti, the ancient DNA data revealed a previously unknown aspect of social organization at this site. Future research at more sites in Nubia, from both the Christian Period and other eras, will help to further elucidate principles of social organization in ancient Nubia.

**Consanguinity**

We identified Runs of Homozygosity (ROH) within the Kulubnarti Nubian individuals with sufficient coverage using the Python package *hapROH* (https://test.pypi.org/project/hapROH/)[1]. We used 5008 global haplotypes from the 1000 Genomes project haplotype panel[40] as the reference panel and applied this method to ancient individuals with a minimum coverage of 400,000 SNPs (n=9, all from the S cemetery) to identify ROH longer than 4 centiMorgan (cM) in the pseudo-haploid data. For each individual, we grouped the inferred ROH into length categories >4cM and >20cM. Large sums of long ROH (>20cM) evidence a close degree of relatedness of the target individual's parents (up to five generations ago), as recombination quickly breaks up blocks back in time, making this signal independent of demographic processes occurring in the deeper past. In contrast, an abundance of shorter ROH signals background parental relatedness and restricted mating pools. We report the total sum ROH in these length bins for each individual with sufficient coverage in Supplementary Table 3 and visualize the size and amount of ROH for all individuals in Supplementary Fig. 2.

Nubia currently has a relatively high rate of consanguinity, characterized by double-first cousin, first cousin, and second cousin marriage[41-43]. It has been suggested that unions between close relatives (with several taboos, including brother-sister, uncle-niece, and aunt-nephew) have been common among Egyptians since the time of the Pharaohs, and that cousin marriages were preferred in Nubians as well, as they ensure the patrilineal system of inheritance[42]. However, we find that only a single individual (I6336/S27, a male who died at approximately nine months old) out of the nine analyzed has closely related parents as indicated by a total of ~160cM ROH in blocks >4cM of with over half of it in segments >20cM. In fact, this individual has an ROH block ~60cM on Chromosome 10 (Supplementary Fig. 3). The amount and length distribution of ROH is typical for offspring of first cousins or genetically equivalent related parents (the average is 220cM ROH with random variation around the average value, into which I6336 falls[1]).

Overall, analysis of ROH suggests that the mating pool of the Kulubnarti Nubians was not sufficiently closed to result in a consistently elevated rate of short ROH. Three individuals have no ROH longer than 4cM at all; three more individuals have no short ROH (4-8cM) which would be expected for a long-standing small population. Of note is some intermediate ROH (8-20cM) present in three out of the nine analyzed individuals, suggesting some mating with a lager meta-population. This signal is consistent with our analysis of admixture dates and ancestry proportions that suggests that admixture at Kulubnarti was ongoing throughout the millennium leading up to and into the Christian Period. Furthermore, our detection that exogamous (here, West Eurasian-related) ancestry was disproportionately associated with female ancestors suggests that these connections could have been primarily female-mediated, and that a mobility system of female exogamy in addition to an inheritance system of patrilineal primogeniture might have been in place at Christian Period Kulubnarti.

### Supplementary Note 4. *qpAdm*

We applied *qpAdm*[44] v.1210 from ADMIXTOOLS[45] with the option 'allsnps: NO' to identify the most likely sources of ancestry and proportions of ancestry in the Kulubnarti Nubians as well as for present-day Nubian groups. We interpreted models as fitting the data at $p>0.05$, and used these models to estimate proportions of admixture.

First, we applied *qpAdm* to investigate if the pooled group of Kulubnarti Nubians (excluding outliers) could be modelled as a result of two-way admixture between Nilotic- and West Eurasian-related ancestry. We used Dinka as a proxy for Nilotic-related ancestry and sought to determine the most appropriate proxy source of West Eurasian-related ancestry, testing 21 geographically and temporally differentiated ancient West Eurasian populations (also including the predominantly West Eurasian-related *Egypt_published*) as possible proxy sources. We began by using the 'O9' reference set (present-day Mbuti, Onge, Chukchi, Karaitiana, Papuan, Han and ancient individuals Ust'-Ishim, MA1, and Kostenki14) that has been previously shown to effectively disentangle divergent strains of ancient West Eurasian-related ancestry (initially defined in ref.[46]; used also in ref.[47]). Results are in Supplementary Data 7.

Upon identifying three plausible solutions for model fit relative to the O9 reference set ($p>0.05$), we implemented a "model competition" approach where a group identified as a possible source relative to the O9 reference set is moved to the reference set if it is not currently being used as a source[46,48]. With this approach, we obtain a fitting model only when *Egypt_published* is used as the West Eurasian-related proxy, but evaluate this as a non-ideal source for accurately estimating ancestry proportions in the Kulubnarti Nubians due to a non-trivial amount of Dinka-related ancestry also present in *Egypt_published* (see Supplementary Data 7 for admixture proportions in *Egypt_published*); in addition, despite the majority proportion of West Eurasian-ancestry in *Egypt_published*, its geographic location reveals that while it is likely to be the proximal source of West Eurasian-related ancestry at Kulubnarti it is not the distal source of such ancestry. For these reasons, we removed *Egypt_published* from our modelling and included the remaining two possible distal sources located in West Eurasia in our model competition approach. We repeated *qpAdm* until all but one admixture model was eliminated; this fitting model used Dinka and *Levant_BAIA* as fitting proxy source to model the Kulubnarti Nubians relative to a reference set that included *Anatolia_EBA* in addition to the O9 reference set populations.

Next, we estimated the proportions of Nilotic- and West Eurasian-related ancestry in each cemetery group (*Kulubnarti_R* and *Kulubnarti_S*) as well as each individual at Kulubnarti using this single fitting admixture model; results are presented in Supplementary Data 7 and Supplementary Data 8. We also used *qpAdm* to determine whether this same admixture model fit three present-day Nubian populations, but found that this model was a poor fit for all present-day targets (Supplementary Data 7).

### Supplementary Note 5. Mitochondrial DNA analysis and haplogroup calling

We determined mitochondrial (mtDNA) haplogroup for each individual in our dataset. We constructed a consensus sequence with samtools v.1.3.1. and bcftools v.1.10.2[49] using a majority rule and aligned mtDNA capture bam files to the *RSRS*[50], restricting to reads with MAPQ ≥ 30 and base quality ≥ 20 and trimming two base pairs to remove deamination artifacts. Haplogroup calls were made using Haplogrep[51] Classify v.2.2.8 with the --rsrs flag. All haplogroups were then assessed as either primarily African- or West Eurasian-associated based on their geographic origin and primary distribution; haplogroup calls, corresponding mutations, and broad geographic groupings provided in Supplementary Data 12, with haplogroups and geographic groupings also depicted in Supplementary Fig. 5. We consider all uniparental data to supplement genome-wide data, which is more broadly informative of an individual's ancestry as it encompasses information from thousands of an individual's ancestors. Here we briefly discuss our mtDNA haplogroup findings; all references to mutations are based on PhyloTree[52] Build 17 with a focus on mutations included in the *RSRS* phylogeny[50].

We found that 35 out of 63 individuals who were not first-degree relatives sharing a maternal lineage belonged to mtDNA haplogroups that originated and are presently distributed predominantly in West Eurasia, while the remaining 28 belonged to African associated haplogroups (all from macrohaplogroup L). In present-day populations, there is a decreased frequency of African mtDNA lineages (L lineages) with a south to north direction[53]; our finding that West Eurasian-associated lineages comprise the slight majority of mtDNA haplogroups at Kulubnarti is consistent with this previous work. There were seven different African-associated mtDNA haplogroups called for the 28 individuals with African-associated haplogroups at Kulubnarti.

Five individuals (three from the R cemetery and two from the S cemetery) belonged to haplogroup L0a1a1, all exhibiting the haplogroup's diagnostic 2759C mutation. Four out of the five individuals (two from the R cemetery and two from the S cemetery) show the same present mutations and three missing mutations, while one of these missing mutations (200G) is found to be present in the remaining individual (I19139/R103). All five individuals have an extra mutation relative to *RSRS* (8017T), while four of these five also have a second mutation (12738C), and two of these four have a third (9102T). L0a1a1 is a subclade of L0a that is estimated to have arisen

~13,500 years ago in eastern Africa[53-55]. The presence of this haplogroup represents deep matrilineal ties to eastern Africa.

Three individuals (all from the S cemetery) belonged to L1b1a2a, with two individuals showing identical haplotypes. All individuals had the 16289G mutation that is diagnostic of this haplogroup and all had an extra mutation at 3357A that was not used in the haplogroup call. In addition, the individual with a different haplotype (I6334/S198) also showed extra mutations at 622T, 10073T, and 12483A. While the parent L1b haplogroup likely arose in West Africa where it is most frequent and diverse, the L1b1a2a lineage likely originated later in East Africa, where it is represented by three divergent sequences from Ethiopia[56]. Previous work has suggested that this haplogroup could have moved from East Africa toward Egypt (where it is also identified) down the Nile River[56], a scenario consistent with its presence at Kulubnarti. No published sequences from Egypt or Ethiopia, or a Bedouin sequence from Israel[57] belonging to haplogroup L1b1a2a show the same unique extra mutations as those found among the L1b1a2a individuals from Kulubnarti.

Fifteen individuals (13 of them not first-degree relatives sharing an mtDNA lineage, three from the R cemetery and 10 from the S cemetery) belonged to haplogroup L2a1d1, the most common mtDNA haplogroup at Kulubnarti. All individuals had the same mutations missing and present; among those present were the five mutations diagnostic of this haplogroup. In addition, all individuals had the extra mutations of 189G, 1872C, 7444A, and 14569A, while a single individual (I6332/S159) also had the extra mutation 6261A. L2a1d1 is an Eastern African subclade of L2a1d that split from L2a1d2 ~10,600 years ago[58]. This haplogroup also represents a deep matrilineal connection to this region[58]. One individual from the R cemetery belonged to another L2a1 lineage, L2a1+143A+16189T (16192T), a branch of L2a1 that likely originated in East Africa. This branch has also been identified in some present-day from the Arabian Peninsula and the Levant in the L2 phylogeny, supporting a long history of gene flow between East Africa and parts of West Eurasia[54].

One individual from the R cemetery belonged to L3b1a2, exhibiting the diagnostic 9300A mutation. This lineage's parent haplogroup (L3b1) is more widespread in Central and West Africa[59], but L3b1a2 has been previously identified in present-day individuals in Egypt[60], Sudan[61] and Somalia[59], suggesting that it is present in also in more easterly parts of Africa, although likely at low frequencies. One individual from the S cemetery belonged to L3f1a1, harboring the five mutations diagnostic of this haplogroup. Haplogroup L3f most likely arose in East Africa; it is most frequent and most diverse in this region[53,59]. L3f1a is one of two main subclades of L3f1, and it is likely that this subclade originated in East Africa, where it is presently found in Somalia and Sudan[59,61]. The haplogroups from the L3 lineage identified at Kulubnarti again represent deep matrilineal ties to East Africa.

Four individuals (two from the R cemetery and two from the S cemetery) belonged to L5a1b; all individuals had the diagnostic 14668T and 14819C mutations and identical haplotypes. The rare L5 haplogroup has been observed at only low frequencies in eastern and into central Africa, including in Egypt, Sudan, Ethiopia, Kenya, Rwanda and Tanzania, as well as in the Mbuti Pygmies[62-68]. L5a1b is estimated to have arisen in East Africa 5,900-15,200 years ago[50], and is today found primarily in Eastern Nilotic speakers[69]. Previous work[57] has identified a present-day Ethiopian as well as an individual from the Sara people of Chad as belonging to L5a1b; these individuals both harbored a TTC insertion between 456-459, which is missing in the four Kulubnarti Nubians belonging to this haplogroup. A Pastoral Neolithic individual from Hyrax Hill in Kenya dating to ~2300 years BP was also assessed as belonging to L5a1b[70]. Consistent with the other L lineages, this haplogroup reflects deep matrilineal ties to East Africa.

In addition to these African-associated mtDNA haplogroups, there were 11 different West Eurasian-associated mtDNA haplogroups represented in 35 individuals at Kulubnarti. While we consider these haplogroups to be West Eurasian-associated given their geographic origin, they were present in northeastern Africa (not only limited to the Nile Valley) possibly for thousands of years before the Christian Period[70-75]. For this reason, it is not possible to assess West Eurasian-related ancestry based on mtDNA haplogroup alone; instead, mitochondrial DNA can be used as a tool for exploring the deep matrilineal origins of the people living at Christian Period Kulubnarti and for investigating patterns of haplotype sharing within and among groups.

mtDNA haplogroup U has a predominant West/Central Eurasian geographic range with branches that also extend into Europe, the Near East, and North Africa[76]. The presence of haplogroup U lineages at Kulubnarti ultimately reflects the biological connections between West Eurasia and Egypt and Nubia established long before the Christian Period. One individual from the R cemetery belonged to U1a1, exhibiting the three diagnostic mutations of this haplogroup but also showing a number of extra mutations to *RSRS*, including 3865G, 6060G, 8544T, 10619T, 14980T, 16183C, and 16319A. U1a1 is predominantly found throughout the Near East and Caucasus, including in people from Yemen, Turkey, and Georgia[77,78]; U1a1 has also been identified in an ancient individual from Egypt dating ~350-200 calBCE[79]. Haplogroup U3, although also primarily found in the Near East and Caucasus, is another one of the branches of macrohaplogroup U that also has a presence in Africa[76]. U3 has been found at relatively low frequencies in present-day Egyptians, Nubians, and Nile Valley groups[62,66]. One individual from the R cemetery belonged to haplogroup U3b, exhibiting all four diagnostic mutations, and harboring a number of extra mutations including 464C, 2272T, 8526C, 9305A, 10909C, 11137C, and 16104T. Haplogroup U3b has previously been identified in geographic proximity to Nubia; specifically, it was called for two ancient Egyptians dating ~750-500 calBCE and ~45 calBCE – 5 calCE[79]. Both U1a1 and U3b have also been previously identified in Bronze Age individuals from Israel and Jordan[80], providing possible evidence of an ancient matrilineal connection to the people living in the Levant, consistent with the genome-wide data reported in this work.

Ten individuals (seven from the S cemetery and three from the R cemetery) belong to haplogroup U5b2b5. All individuals have the same three missing mutations, the same mutations present (including the two mutations diagnostic of this haplogroup), and the same three extra mutations (13980A, 15226G, 15538T); this suggests that there is plausibly an unidentified sub-lineage of U5b2b5. U5 is known for being one of the most ancient mtDNA haplogroups in Europe[81,82], primarily identified in Mesolithic hunter-gatherers[83,84], and U5b2 has been shown to be the most ancient sub-haplogroup of U5b[85]. Haplogroup U5b2b5 was called for a 4,000-year-old mummy from Egypt[86]; this individual also shared the extra 15538T mutation called for the Kulubnarti Nubians, suggesting that it is possible that this haplogroup was spread into Nubia via Egypt.

One individual belonged to T1a7, exhibiting the three diagnostic mutations of this haplogroup and also exhibiting extra mutations including 319C, 5201C, 5460A, and 16172C. Haplogroup T has an unambiguous Near Eastern origin[81], and some of its lineages have been found at different frequencies in populations throughout Egypt and the Nile Valley[60,62,87]. The T1a lineage split ~17,000 years ago, and several individuals from Egypt[60] as well as throughout the Near East, including Israel and Iraq[88], have been shown to have the mutations consistent with T1a7; more recently, present-day individuals from Lebanon have been shown to belong to T1a7[89]. Possibly of direct relevance is the detection of the T1a7 lineage in ancient Egypt dating ~800 BCE–1 CE[79,90], which plausibly could have been spread southward into Nubia prior to the Christian Period.

One individual from the R cemetery belonged to R0a1a, harboring the four diagnostic mutations of this haplogroup as well as additional mutations at 8527G, 9631C, 11167G, and 15779C. Haplogroup R0a is most frequent in the Arabian Peninsula and Horn of Africa; previous work has proposed that the deep presence of R0a in Arabia highlights at least one Pleistocene glacial refugium on the Red Sea plains, and that the dispersal of this haplogroup into East Africa occurred at the end of the Late Glacial[91], giving this haplogroup a deep presence in Africa as well as on the Arabian Peninsula (as well as throughout other parts of West Eurasia). In particular, R0a1a is one of the known major expansion lineages in R0a; however, the vast majority of African R0a lineages fall within R0a2[91]. While R0a1a is more represented on the Arabian Peninsula[92], it has previously been identified in Egypt, including in two individuals reported in ref.[79], one dating ~360–210 BCE and the other ~40 BCE–15 CE. The presence of this lineage at Kulubnarti likely represents deep connections between the Arabian Peninsula and East Africa that resulted in genetic exchange long before the Christian Period.

One individual from the S cemetery belonged to N1a1a. This haplogroup has eight diagnostic mutations, and this individual harbored seven of them, missing the 16147G transversion. Additional mutations included 10586A, 10768G, 13146T, 16147A, and 16245T. N1a originated in the Near East[81] and is now widely distributed across the Near East, Europe, northeast Africa, and into Central Asia. N1a displays deep diversity in eastern Africa as well as the southern part of the Arabian Peninsula and probably reflects ancient gene flow (plausibly dating to the Late Glacial period)[93]; in northeast Africa, the N1a haplogroup is primarily found in groups speaking Afro-Asiatic languages who have substantial amounts of ancestry with an ultimate origin in West Eurasia. Haplogroup N1a1a in the Horn of Africa is believed to have also spread in the Late Glacial[93], suggesting its presence in northeast Africa for thousands of years before the Christian Period.

Three individuals (two from the S cemetery and one from the R cemetery) belonged to N1b1a2. All exhibited the diagnostic 4904T mutation, and the two individuals from the S cemetery shared the same haplotype. In modern populations, haplogroup N1b1 is found primarily in the Near East, with branches in Europe and North Africa[93]. Haplogroup N1b1a has been previously identified at the Anatolian Ceramic Neolithic site of Barcin (6500-6200 BCE)[94], while N1b1a2 has been previously found in Bronze Age Israel and Jordan[80], again providing evidence of a West Eurasian-associated matrilineal connection as also shown through genome-wide data.

One individual from the R cemetery belonged to K1a19, harboring the 12338C mutation diagnostic of this haplogroup and additional mutations including 5563A and 15929G. Haplogroup K is most often associated with Neolithic farmers[95]; it spread and diversified during the Neolithic expansion into Europe, and it is also found in Central Asia and in the Horn of Africa. K1a19 is a rare haplogroup[96] that is reported to have origins in the Near East, though it is also spread throughout other regions, including southern Europe and Iran[97].

One individual from the S cemetery belonged to HV13a, although this individual did not have one of the seven diagnostic mutations of this haplogroup (9027T); an additional mutation at 8420G was also observed. While haplogroup HV has a likely Near Eastern origin (Derenko et al. 2014), it was detected at ~14% frequency in a small population from the Egyptian Western Desert (west of the Nile River), providing direct genetic evidence of a strong Near Eastern genetic input into this region that dates to the Neolithic[60]. Haplogroup c has been shown to have a Near Eastern origin[98]; to our knowledge, this specific lineage has not been detected in Africa.

Two individuals (one from the R cemetery and one from the S cemetery) belonged to haplogroup J2a2e, harboring both of the diagnostic mutations (10658A, 14364A) of this haplogroup. These individuals have the same haplotype, which includes the additional mutations 9276A, 14016A, and 16362C. Haplogroup J2 originated in the Near East[88], with the J2a lineage estimated to be between ~20,000 and 28,000 years old[50]. The J2a2e haplogroup was also called for two ancient Egyptian individuals reported in ref.[79] (also see ref.[90]), one dating to ~350–200 cal BCE and the other to ~80–130 CE, while a present-day Egyptian individual belonging to J2a2e was also shown to have the same three additional mutations as seen in the Kulubnarti Nubians[99]. As such, while this haplogroup originates in the Near East, it is more likely to reflect biological connections between Nubia and Egypt.

Fourteen individuals (eight from the R cemetery and six from the S cemetery, 13 who were not first-degree relatives sharing a mtDNA lineage) were assigned as belonging to haplogroup H2a. To our knowledge, haplogroup H2a has not previously been found in any ancient African individuals. Macrohaplogroup H is the predominant West Eurasian haplogroup that comprises nearly a half of the European mtDNA pool and decreases to frequencies ~10-30% in the Near East and Caucasus[81]; H2a is one of the sub-haplogroups of this lineage that exhibits a distinct phylogeographic pattern[100]. The spread of H2a extends to Central Asia[100] and it is more often associated with eastern European affinity than western European affinity[101-103]. Evidence for H2a in Africa is sparse: it has been reported for a small number of Tunisian Berbers and other North Africans[104], but otherwise appears to be largely absent in African individuals.

All individuals assigned as belonging to H2a had the diagnostic 4769A mutation for this haplogroup and all had three additional mutations (15784C, 16210G, and 16224C). Particularly interesting are the latter two additional mutations: 16210G is not reported in PhyloTree 17, while 16224C is part of PhyloTree 17 but is not part of the H2a haplogroup. This raises the possibility that these individuals were erroneously assigned to H2a based on the presently-available version of PhyloTree, but were actually part of a now extinct or previously undocumented mtDNA haplogroup.
